# Unilateral acoustic degradation delays attentional separation of competing speech

**DOI:** 10.1101/2020.11.20.391391

**Authors:** Frauke Kraus, Sarah Tune, Anna Ruhe, Jonas Obleser, Malte Wöstmann

**Affiliations:** Department of Psychology, University of Lübeck, Lübeck, Germany; Center of Brain, Behavior and Metabolism (CBBM), University of Lübeck, Lübeck, Germany

**Keywords:** speech tracking, envelope, unilateral vocoding, cochlear implant, attention, spectro-temporal response function

## Abstract

Hearing loss is often asymmetric, such that hearing thresholds differ substantially between the two ears. The extreme case of such asymmetric hearing is single-sided deafness. A unilateral cochlear implant (CI) on the more severely impaired ear is an effective treatment to restore hearing. The interactive effects of unilateral acoustic degradation and spatial attention to one sound source in multi-talker situations are at present unclear. Here, we simulated some features of listening with a unilateral CI in young, normal-hearing listeners (*N* = 22) who were presented with 8-band noise-vocoded speech to one ear and intact speech to the other ear. Neural responses were recorded in the electroencephalogram (EEG) to obtain the spectro-temporal response function (sTRF) to speech. Listeners made more mistakes when answering questions about vocoded (versus intact) attended speech. At the neural level, we asked how unilateral acoustic degradation would impact the attention-induced amplification of tracking target versus distracting speech. Interestingly, unilateral degradation did not per se reduce the attention-induced amplification but instead delayed it in time: Speech encoding accuracy, modelled on the basis of the sTRF, was significantly enhanced for attended versus ignored intact speech at earlier neural response latencies (<~250 ms). This attentional enhancement was not absent but delayed for vocoded speech. These findings suggest that attentional selection of unilateral, degraded speech is feasible, but induces delayed neural separation of competing speech, which might explain listening challenges experienced by unilateral CI users.

## Introduction

Selecting the relevant auditory signal from a mixture of sounds is a challenging task. Selective attention enables the listener to prioritize target information against concurrent distraction. In recent years, methods have been developed to quantify the phase-locking of neural responses in the magneto/electroencephalogram (M/EEG) to the envelope of individual speech signals in the mixture of sounds (for reviews, see Brodbeck & Simon, 2020; Ding & Simon, 2014; Peelle & Davis, 2012; Wöstmann, Fiedler, et al., 2017). A number of studies have convergingly shown that normal-hearing listeners’ M/EEG responses exhibit stronger neural phase-locking to the envelope of attended versus ignored speech (e.g. Ding & Simon, 2014; Horton et al., 2013; O’Sullivan et al., 2015; Zion Golumbic et al., 2013). Furthermore, there is evidence that the degree of neural phase-locking to the envelope of speech correlates with speech intelligibility (e.g. Peelle et al., 2013) and with behavioural indices of speech comprehension (e.g. Etard & Reichenbach, 2019). The neural phase-locking to the speech envelope – also referred to as “neural speech tracking” (Obleser & Kayser, 2019) – constitutes an objective measure of the attentional enhancement of target speech against distraction.

Important for the present study, research has associated auditory processing deficits with particular changes in neural speech tracking. For instance, the differential cortical tracking of attended versus ignored speech was found to be reduced in listeners with stronger hearing loss in some studies (Petersen et al., 2017), although results of other studies suggest stronger tracking of target speech with increasing hearing loss (Decruy et al., 2020; Fuglsang et al., 2020). Furthermore, noise-vocoding to simulate hearing with a cochlear implant (CI) reduced the attentional enhancement of target speech (Ding et al., 2013; Kong et al., 2015; Rimmele et al., 2015). A recent study found that bilateral CI users showed smaller early neural separation of attended versus ignored speech (<200 ms) but larger late neural separation (>200 ms) compared with normal-hearing controls (Paul et al., 2020). These findings demonstrate the potential of neural speech tracking to understand interactive effects of degraded acoustics and attention.

A so far neglected test case for the impact of acoustic degradation on neural speech tracking is asymmetric hearing. There are at least three reasons why asymmetric hearing poses a relevant test case for neural speech tracking. First, asymmetric hearing provides a well-controlled scenario to study the simultaneous neural tracking of degraded versus intact acoustic input in the same neural system. In contrast, comparing speech tracking in hearing-impaired versus normal-hearing listeners poses the challenge to control for between-subject differences. In such cases, it often remains somewhat unclear to what extent changes in speech tracking reflect differences in the perceived acoustic input versus differences in neural processing capabilities. Second, the prevalence of asymmetric hearing loss in the population should not be disregarded (e.g. Golub et al., 2018; Wilson et al., 1993), which emphasizes the importance to study auditory selective attention in such cases. Third, although unilateral CI implantation can partially restore hearing (for review, see Vlastarakos et al., 2014), effects of the CI-induced unilateral spectral degradation on the attentional selection of target speech against concurrent acoustic masking are largely unknown.

Noise vocoding is a common technique used to simulate some aspects of hearing with a CI. In essence, noise vocoding degrades the spectral information of the acoustic input but leaves the temporal envelope largely intact (Rosen et al., 1999). When applied to target speech, noise vocoding compromises speech comprehension in silence (Shannon et al., 1995) and particularly in background noise (for review, see Moore, 2008). Noise-vocoding of speech distractors has been shown to reduce the extent of distraction, evidenced by better target speech reception (e.g. Westermann & Buchholz, 2017) and better memory for target speech (e.g. Ellermeier et al., 2015; Senan et al., 2018; Wöstmann & Obleser, 2016).

Noise vocoding affects the type of masking elicited by speech distractors: Energetic (acoustic) masking is complemented by informational masking for vocoded speech signals that are still intelligible (e.g. Brungart et al., 2001; Mattys et al., 2009), for instance through noise vocoding with a larger number of vocoder bands. Furthermore, it has been proposed that informational masking additionally increases with higher similarity of target and distracting speech (e.g. Rosen et al., 2013). In consequence, when competing speech signals are presented dichotically, a unilateral CI user should experience a relatively higher degree of informational masking when the masking speech distractor is presented to the better ear and is thus more intelligible.

Different modulations of neural speech tracking are conceivable that might accompany the attentional selection of a unilateral speech signal that is acoustically degraded versus intact. It has been proposed that auditory selective attention depends on two closely intertwined processes, referred to as *auditory object formation* (grouping of sounds that originate from the same physical source) and *auditory object selection* (selecting an object for prioritized processing at the expense of distraction; Shinn-Cunningham, 2008). The neural tracking of speech, which reflects both bottom-up acoustic information such as the syllabic structure as well as top-down attention, might be a neural correlate of both, auditory object formation and selection (Shinn-Cunningham et al., 2017). We have previously observed that earlier components of the speech tracking response are mainly sensitive to acoustic features, whereas later response components reflect the attention focus and the effort of attentional selection (Fiedler et al., 2019). In agreement with this, a recent study proposed a model wherein earlier neural tracking responses (< ~100 ms) reflect filtering and restoration of acoustic signals, whereas later responses correspond to active attentional streaming (Brodbeck et al., 2020). Thus, the latency at which unilateral acoustic degradation impacts neural speech tracking in the present study allows to draw conclusions about the underlying perceptual and cognitive mechanisms.

The present study implemented a dichotic listening task to present an intact speech signal to one ear and a different, noise-vocoded speech signal to the other ear. This way, the present study deliberately avoided simulation of listening challenges associated with the integration of acoustic hearing (with the non-implanted ear) and electric hearing (with the implanted ear) of sound emitted from a single physical source (see e.g. Litovsky et al., 2019; Ma et al., 2016). Spatial cues guided listeners’ attention either to the intact or noise-vocoded target speech, while the respective other speech signal served as a to-be-ignored distractor. Since noise vocoding affects the spectral integrity of acoustic signals, we used EEG recordings to derive temporally-but also spectrally-resolved neural response functions. We predicted an attention-induced increase in the neural response to attended compared to ignored speech, which would decrease for attending-versus-ignoring vocoded compared to intact speech. It was, however, an open question at which neural response latency unilateral acoustic degradation would affect the tracking of the speech envelope.

## Materials and Methods

### Participants

We tested N = 22 young, normal-hearing adults (age: 19-33 years; mean = 24.6 years; SD = 3.76 years; 12 females; all right-handed). Pure tone audiometry ensured that the pure tone average (PTA; average hearing threshold across frequencies 500, 1,000, 2,000, and 4,000 Hz) per ear was below 20 dB HL (hearing level). The mean PTA on the right side was 5.18 HL (SD = 5.97 dB) and on the left side 2.68 HL (SD = 3.59 dB). Two participants had a moderate hearing loss of up to 40 dB HL at the frequencies 500, 750 and 1,000 Hz on the right side. Due to the overall good behavioural performance of those participants and PTA smaller than 20 dB HL, we decided to not exclude the data of these participants. Participants were recruited via the participant database of the Department of Psychology at the University of Lübeck. They were native speakers of German and reported no history of neural disorders.

Participants gave their written informed consent and were financially compensated with €10/hour or received course credit. The study was approved by the local ethics committee of the University of Lübeck.

### Stimuli

Two German audiobooks were used as continuous speech streams (“Eine kurze Geschichte der Menschheit” by Yuval Noah Harar, spoken by a male narrator, “Nero Corleone kehrt zurück - Es ist immer genug Liebe da” by Elke Heidenreich, spoken by a female narrator). Of the first 64 minutes of each audiobook, we used the first half for the to-be-attended stream, and the second half for the to-be-ignored stream. We generated individual trials by cutting the respective portions of the audiobook into segments of four minutes each.

For noise-vocoding, we used a gamma tone filterbank (implemented in the Auditory Toolbox for Matlab; Slaney, 1993) with eight logarithmically-spaced filters between 0 and 8 kHz. After filtering the speech stimuli with these filters, we multiplied the envelope (magnitude of the Hilbert transform) in each frequency band with white noise filtered with the same filterbank and applied a root mean square (RMS) normalization. Finally, the eight bands were summed and again normalized to the RMS value of the intact audiobook parts (Shannon et al., 1995). We used vocoding with 8 frequency bands to balance task difficulty in a way that participants could perform the task well above chance level but that listening to vocoded speech was more challenging than listening to intact speech.

### Procedure

Participants performed the continuous speech task sitting in a sound-attenuating chamber. The audiobooks were presented dichotically through in-ear headphones (EARTONE 3A, 3M). Participants could set the overall volume to a comfortable level before the experiment started. During the whole experiment the input to one ear was vocoded speech while the input to the other was intact speech. Therefore, there were two experimental settings. In setting 1, participants attended to the intact speech and ignored the vocoded distractor stream. By contrast, in setting 2, they attended to the vocoded speech and ignored the intact speech stream. The side of vocoding was counterbalanced across participants.

In total, the experiment consisted of 16 trials with 4 minutes duration each. Each of the two experimental settings was presented in 8 trials of the experiment. Participants started with two attention-to-intact speech trials, followed by two attention-to-vocoded speech trials and so on. The to-be-attended audiobook (male versus female talker) alternated from each trial to the next. Before each trial, a spatial cue and the text “Listen to the left/right” signalled the to-be-attended side and participants started the trial with a button press on a four-button response box (The Black Box Toolkit, Sheffield, UK). After each trial, participants answered four multiple choice questions, each one with four alternative response options about the content of the to-be-attended audiobook. During the whole task we measured participants’ electroencephalogram (EEG).

### EEG recording and preprocessing

The EEG was recorded from 64 electrodes (ActiChamp, Brain Products, München) at a sampling rate of 1,000 Hz, referenced to electrode Fz. After recording, the EEG data were preprocessed with the FieldTrip toolbox (version 2019-09-20; Oostenveld et al., 2011) for Matlab (MathWorks, Inc.). The data were re-referenced to the average of all electrodes and filtered between 1 Hz (high-pass) and 100 Hz (low-pass). Following independent component analysis (ICA), components corresponding to eye blinks, lateral eye movements, muscle activity and heartbeat were rejected upon visual inspection. The cleaned data were low-pass filtered at 10 Hz, cut into trials of four minutes starting at the onset of the auditory stimulus and resampled to 125 Hz. For the subsequent analysis these trials were cut into four consecutive one-minute segments and z-transformed (Crosse et al., 2016).

### Extraction of stimulus features

The experiment was designed to simulate some features of listening with a unilateral CI. We used the onset envelopes of the intact speech signal to estimate the spectro-temporal response function (sTRF, see below). The NSL toolbox (Chi et al., 2005) for Matlab was used to compute a spectrally resolved representation of the intact speech stimuli. We extracted an auditory spectrogram consisting of 128 logarithmically spaced frequency bands. To extract the onset envelope in each frequency band, we computed the envelope (magnitude of the Hilbert transform), took the first derivative, and applied half-wave rectification. After down-sampling in the time and frequency domain, the envelopes of the auditory stimuli included 16 frequency bands (logarithmically spaced between 0 and 8 kHz) and had the same sampling frequency fs = 125 Hz as the EEG data. A spectral resolution of 16 frequency bands for the sTRF analysis was used to yield a spectral resolution higher than the number of vocoder bands but still low enough to avoid redundancy.

The major reason for using the onset envelope instead of the conventional (unprocessed) envelope of the speech signal for sTRF modelling was that previous studies (Drennan & Lalor, 2019; Fiedler et al., 2017) found that emphasizing the onsets in the envelope increased the neural encoding response. These observations speak to the view that the brain tracks local changes in amplitude (“edges”) rather than the absolute amplitude of the speech envelope (Oganian & Chang, 2019). Therefore, we used the onset envelope to induce a larger and more robust speech tracking response, which should benefit the statistical power to observe modulatory effects of acoustic detail and attention.

### sTRF estimation

The sTRF was estimated using the mTRF Toolbox (Crosse et al., 2016) for Matlab using regularized linear regression. First, for every one-minute segment, a forward (‘encoding’) model m was estimated with *m* = (*S^T^S* + *λI*)^−1^*S^T^R*. S is a matrix containing the onset envelopes of the attended and ignored speech stream and their time-lagged versions of the attended and the ignored speech stream. We used time lags in the range from −150 to +450 ms. The matrix R contains the corresponding EEG signal of all 64 electrodes, and I is the identity matrix. Leave-one-out cross-validation was used to determine the optimal λ parameter, which maximized the Pearson correlation between the predicted and measured neural responses. To allow for an unbiased comparison between the different conditions, the output data of this estimation were z-scored. Models of attended and ignored speech that belonged to the same experimental trial were jointly estimated. That is, models of to-be-attended intact speech and to-be-ignored vocoded speech were trained together as were the models of to-be-attended vocoded speech and to-be-ignored intact speech. In total, 32 individual models were trained per participant and condition and averaged. For illustration purposes only, the sTRF was also averaged over nine central electrodes (FC1, FCz, FC2, C1, Cz, C2, CP1, CPz, CP2).

### Neural encoding accuracy

Further, the sTRF was used to predict the EEG response to a given speech segment. To this end, using a leave-one-out approach, we convolved the envelopes with the trained sTRF models averaged across all but the current segment. Afterwards, we compared each predicted EEG response to the empirical EEG response using Pearson correlation to assess how strongly a speech signal is encoded by the neural response. The resultant r values were averaged per participant and condition and reflect a continuous, fine-grained measure of neural tracking encoding accuracy (Fiedler et al., 2017; Fiedler et al., 2019; Fuglsang et al., 2020).

To analyse the temporal unfolding of neural encoding accuracy across sTRF time lags, we predicted the EEG response based on a 48-ms sliding window (24 ms overlap) of the sTRF (Fiedler et al., 2019; Jessen et al., 2019). In essence, we used each 48-ms time window of the sTRF separately to predict the EEG-Signal. This approach enabled us to test which time windows of the sTRF contribute strongest to EEG prediction. As before, the calculation was carried out using a leave-one-out procedure. The obtained Pearson correlation coefficients were averaged per time window, condition, and individual participants.

Furthermore, for illustration purpose, we analysed the spectral unfolding of the neural encoding accuracy across the frequency bands of the sTRF. For each frequency band of the sTRF, we used sliding 48-ms time windows to predict the EEG-signal. This resulted in temporal and spectral unfolding of the neural encoding accuracy.

### Statistical Analysis

For behavioural data analysis, we used dependent-samples and independent-samples non-parametric permutation tests on the proportion of correctly answered multiple-choice questions. In detail, a dependent-samples permutation test was used to contrast the proportion of correctly answered content questions about to-be-attended intact versus vocoded speech across all participants (*N* = 22). Two independent-samples permutation tests were used to contrast proportion correct scores for to-be-attended intact speech presented on the left (*n* = 11) versus right side (*n* = 11), and similarly for to-be-attended vocoded speech presented on the left (*n* = 11) versus right side (*n* = 11). Reported *p*-values (*p_permutation_*) correspond to the proportion of absolute values of 5,000 *t*-statistics computed on data with permuted condition labels that exceed the absolute empirical *t*-value for the original (unpermuted) data (Wöstmann et al., 2019). Note that dependent-samples non-parametric permutation tests were furthermore used to contrast neural encoding accuracy derived from the entire sTRF across conditions.

To analyse possible differences between the sTRF in the four conditions of the 2 (acoustic detail: intact vs. vocoded) x 2 (attention: attended vs. ignored) design, we used a cluster permutation test implemented in the FieldTrip toolbox (Maris & Oostenveld, 2007) for Matlab. For this statistical analysis we used the data of all 64 electrodes. First, we calculated two difference-sTRFs for each participant through subtraction of the ignored from the attended sTRF, separately for intact and vocoded speech. The resulting difference-sTRFs [intact (attended–ignored), vocoded (attended–ignored)] were contrasted using multiple dependent-samples *t*-tests for all time-frequency-electrode triplets of the sTRF. Resulting *t*-values were then used to identify clusters by clustering neighbouring time-frequency-samples with *p* < 0.05 for at least three neighbouring electrodes. In each observed cluster, the *t*-values were summed (reported as *t*-sum) over all bins belonging to the cluster (reported as *No. of cluster bins*) and compared to a permutation distribution. The permutation distribution was generated by randomly assigning the sTRF data to conditions, followed by summation of their *t*-values, through 5,000 iterations. In order to visually resolve the interaction, we calculated multiple dependent-samples *t*-tests for all time-frequency-electrode triplets of the sTRF to contrast the attended with the ignored sTRF (i.e., main effect attention), separately for intact and vocoded speech. Significant clusters for the interaction (acoustics x attention) were then overlaid on top of the resulting *t*-maps.

For statistical analysis of temporally-resolved encoding accuracy, we used cluster permutation tests as well. First, two cluster permutation tests were used to contrast encoding accuracy for attended versus ignored speech, separately for intact and vocoded speech. In order to test the acoustics x attention interaction, we used a cluster permutation test to contrast the attended–ignored difference of encoding accuracy for intact versus vocoded speech.

## Results

### Unilateral degradation decreases recall of attended speech content

Participants performed a spatially cued dichotic listening task (Fig. 1A). The assignment of vocoded speech to the left or right ear was counterbalanced across participants. On a given trial, participants were cued to attend to the speech signal on the left or right side. At the end of each trial, participants answered four multiple-choice questions about the content of the to-be-attended audiobook.

**Figure 1.**
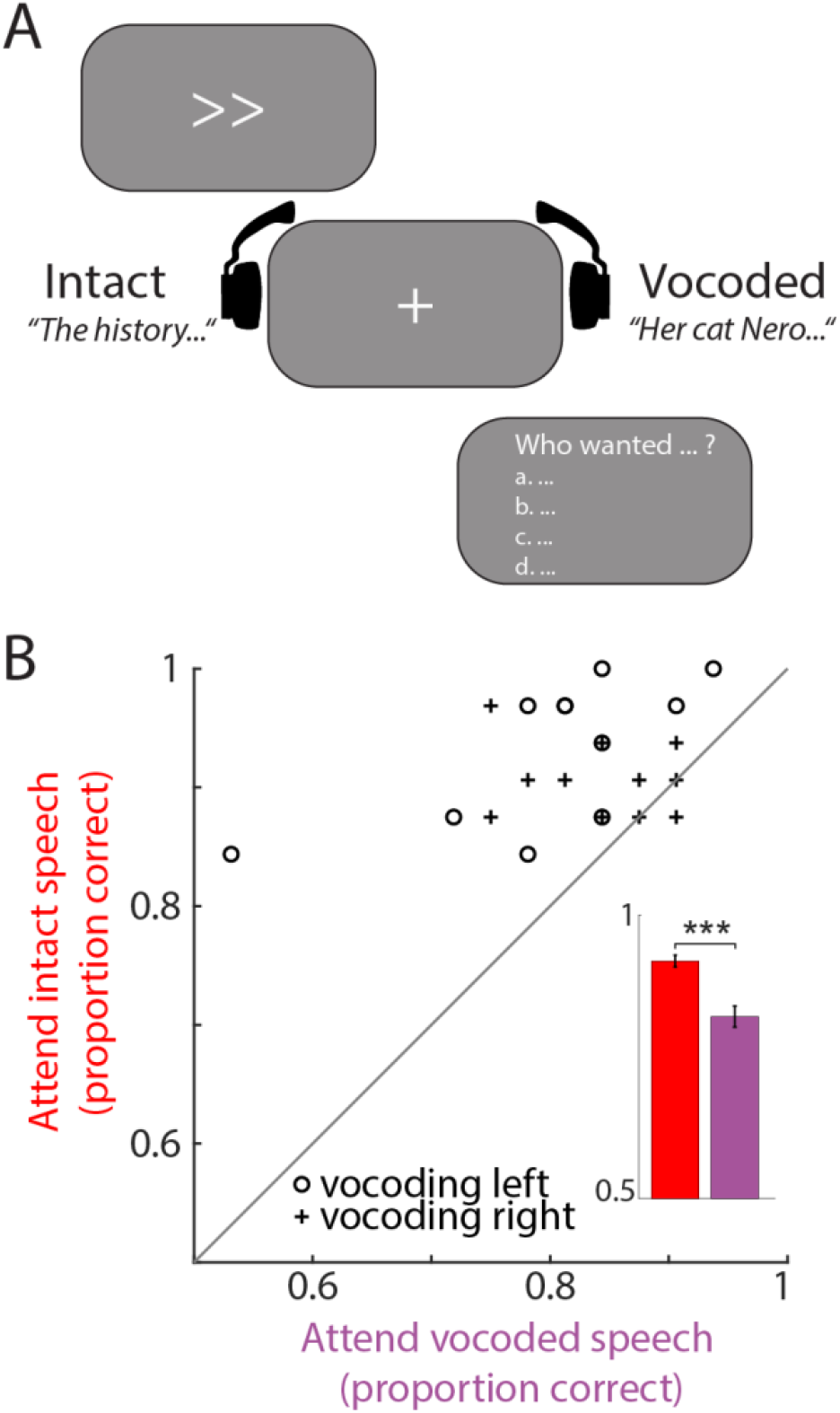
Experimental design and behavioural results. (**A**) Design of continuous speech task. On each trial, a visual spatial cue and the text “Listen to the left/right” guided participants’ attention to the left or right side. During the entire experiment, participants listened to intact speech on one ear and to vocoded speech on the other ear (balanced across participants). After each trial, participants had to answer four closed four-choice questions about the content of the to-be-attended audiobook. (**B**) The 45-degree plot contrasts single-subject proportions of correct answers to content questions for attention to intact versus vocoded speech for participants with vocoded speech on the left side (circles) or right side (crosses). Note that some data points fall on top of each other. Bars and error bars in the inset show mean proportion correct ±1 SEM, respectively. *** *p_permutation_* < 0.001.

Individual participants’ answers to content questions (Fig. 1B) were correct well above chance level (here: proportion correct of 0.25). As expected, the proportion of correct responses to content questions was significantly higher when the to-be-attended audiobook was spectrally intact (mean proportion correct = 0.92; SEM = 0.001) compared to vocoded (mean proportion correct = 0.82; SEM = 0.0018; *p_Permutation_* = 0.0002; Cohen’s d = 1.195; for comparable behavioural performance modulation in a small group of unilateral CI users in a pilot experiment, see Supplementary Materials and Supplementary Fig. S1). Of note, this effect was very consistent across participants (i.e., present in 19 of 22 participants). The side of speech presentation (left for *n* = 11 and right for *n* = 11) did not significantly affect the proportion of correct responses to content questions about intact speech (left vs. right: *p_permutation_* = 0.1722) or vocoded speech (left vs. right: *p_permutation_* = 0.2899).

### Spectro-temporal responses to unilateral intact and vocoded speech

In the sTRF that models the neural impulse response to the speech envelope extracted from different frequency bands of the acoustic signal, we observed the typical morphology, including earlier P1, N1, and P2 response components, as well as a later N2 response component (Fig. 2). Notably, these response components were strongest for speech envelopes extracted from auditory signals below ~1,000 Hz and weaker for higher frequencies up to 8,000 Hz. Since we had no prior hypotheses regarding effects of the side of vocoding (left vs. right) and since there are no obvious differences between sides of vocoding in Figure 2, we collapsed across these two conditions and focused all subsequent analyses on the factors acoustic detail (intact vs. vocoded) and attention focus (attended vs. ignored).

**Figure 2.**
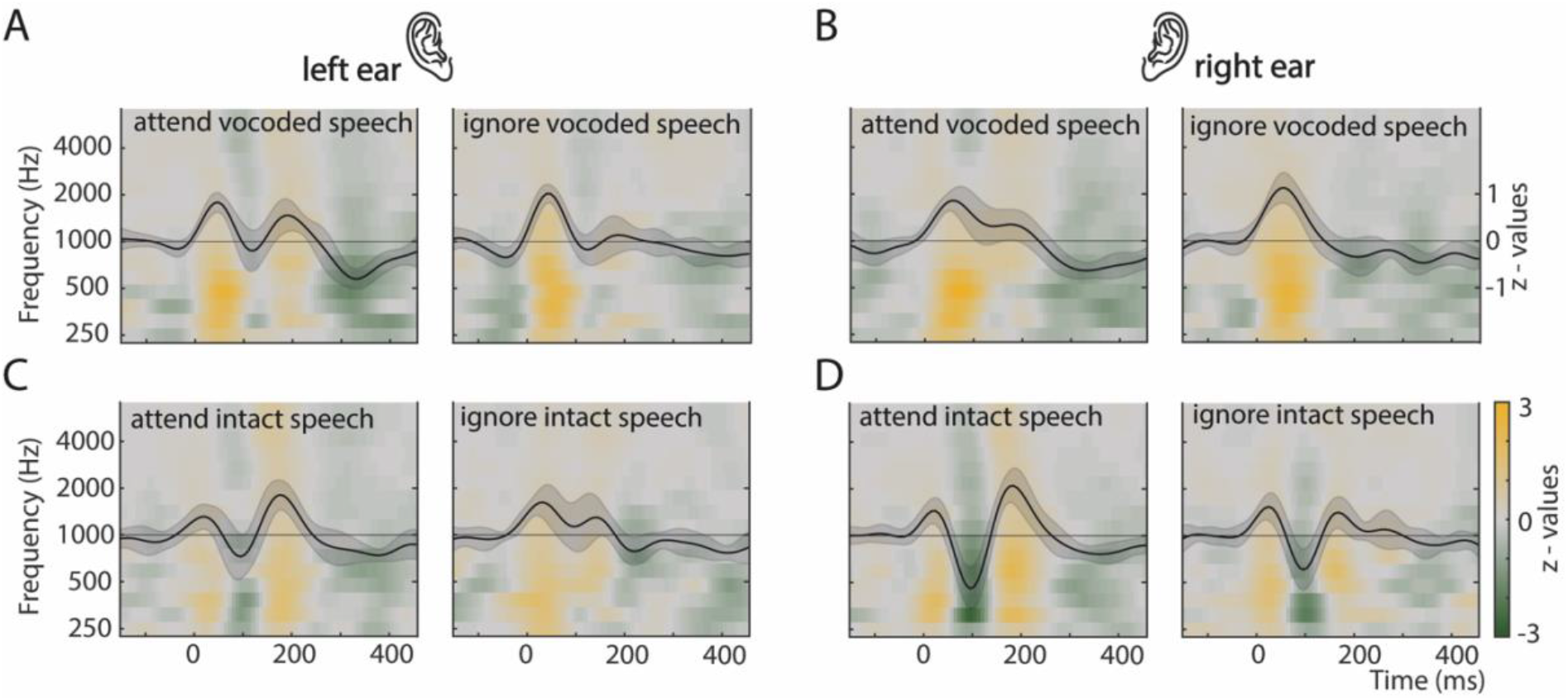
Spectro-temporal response functions for all experimental conditions. Time-frequency panels show average spectro-temporal response functions (sTRF); solid black lines and shaded areas show the temporal response function (TRF; marginalized across frequencies of the sTRF) and 95-% confidence intervals, respectively. The four left-side panels show the response functions for speech materials presented to the left ear split by acoustics and attention condition, the four right-side panels show the same for input to the right ear. Within each participant, neural responses were averaged across nine central electrode positions (see Materials and Methods for details). Note that A & D show data of *n* = 11 participants with vocoded speech presented to the left ear and intact speech to the right ear. C & B show data of *n* = 11 participants with vocoded speech presented to the right ear and intact speech to the left ear.

### Interactive effects of unilateral degradation and attention on neural speech tracking

Figure 3 shows interactive effects of unilateral vocoding and attention on the sTRF (panels A&B), as well as on the marginalized temporal response function (TRF; panels C&D). The corresponding statistical analysis of neural response functions was divided into two steps. First, we analysed the interactive effect of acoustic detail (intact vs. vocoded) x attention focus (attended vs. ignored) on the sTRF (Fig. 3A). The interaction was significant in two fronto-central electrode clusters for frequencies below ~2,000 Hz in time intervals overlapping mainly with the N1 response component (negative cluster with *cluster p-value* = 0.0014, 40-152 ms, 199-1,850 Hz; *t*-sum = −2,104; No. of cluster bins = 703) and with the N2 response component (positive cluster with *cluster p-value* = 0.0002, 176-328 ms, 199-1,850 Hz; *t*-sum = 2,972; No. of cluster bins = 991).

**Figure 3.**
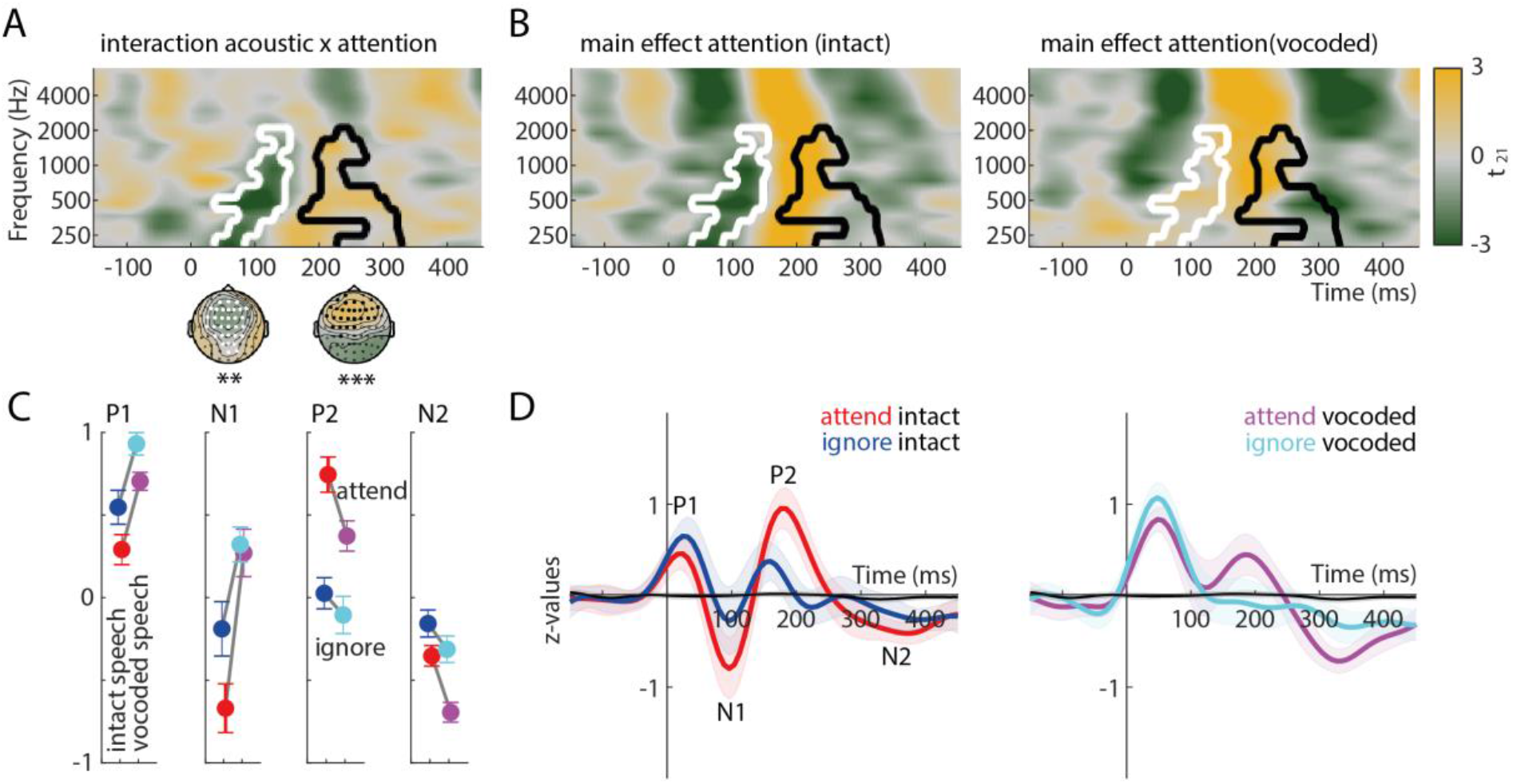
Effects of acoustics and attention on the spectro-temporal response function. (**A**) The time-frequency plot shows *t*-values for the interaction effect of acoustic detail (intact vs. vocoded) x attention (attended vs. ignored) on the spectro-temporal response function, averaged for illustrative purposes across nine central electrodes. White and black outlines indicate a significant negative (*p* = 0.0014 **) and positive cluster (*p* = 0.0002 ***), respectively. Topographic maps show average cluster *t*-values; electrodes belonging to the earlier and later significant cluster are highlighted in white and black, respectively. (**B**) Time-frequency plots shows *t*-values for the main effect of attention (attended vs. ignored) for intact speech (left) and vocoded speech (right), overlaid with outlines of significant acoustics x attention clusters from (A). (**C**) For illustration purposes, we averaged the sTRF across frequencies, nine central electrodes, and distinct time periods corresponding to individual response components (P1: 16-56 ms, N1: 80-120 ms, P2: 176-224 ms, N2: 312-352 ms). Dots and error bars show the average across participants ±1 SEM, respectively. (**D**) Solid lines and shaded areas show average temporal response functions (TRF; marginalized across frequencies of the sTRF) and 95-% confidence intervals.

Second, in order to resolve the significant acoustic detail x attention focus interaction, we analysed the simple main effect of attention focus on the sTRF, separately for intact and vocoded speech signals (Fig. 3B). Interestingly, the earlier interaction cluster was driven by a larger (more negative) N1 component for attended than ignored intact speech, while the same effect was virtually absent for vocoded speech. By contrast, the later interaction cluster was mainly driven by a larger (more negative) N2 component for attended than ignored vocoded speech, while this effect was reduced for intact speech (see also Fig. 3C&D).

In addition to interactive effects of vocoding and attention, it is obvious from Figure 3C&D that vocoded speech elicited a generally larger early P1 response component compared with spectrally intact speech (see also Supplementary Figure S2 for a grouping of the TRF according to conditions occurring at the same time in the experiment: attend intact, ignore vocoded speech; attend vocoded, ignore intact speech).

### Acoustic degradation delays attentional selection of target speech

In a final analysis, we focused on a more fine-grained temporal representation of the attentional enhancement of target versus distracting speech, that goes beyond individual components of the sTRF. To this end, we applied a sliding-window approach to predict the EEG signal at individual electrodes from the sTRF and speech envelopes (Fig. 4). A more positive correlation of the predicted with the measured EEG signal points to a stronger neural representation of the respective speech signal, referred to as higher *neural encoding accuracy*. We asked in which time windows attention would enhance the encoding accuracy of target versus distracting speech signals in case these were intact versus vocoded.

**Figure 4.**
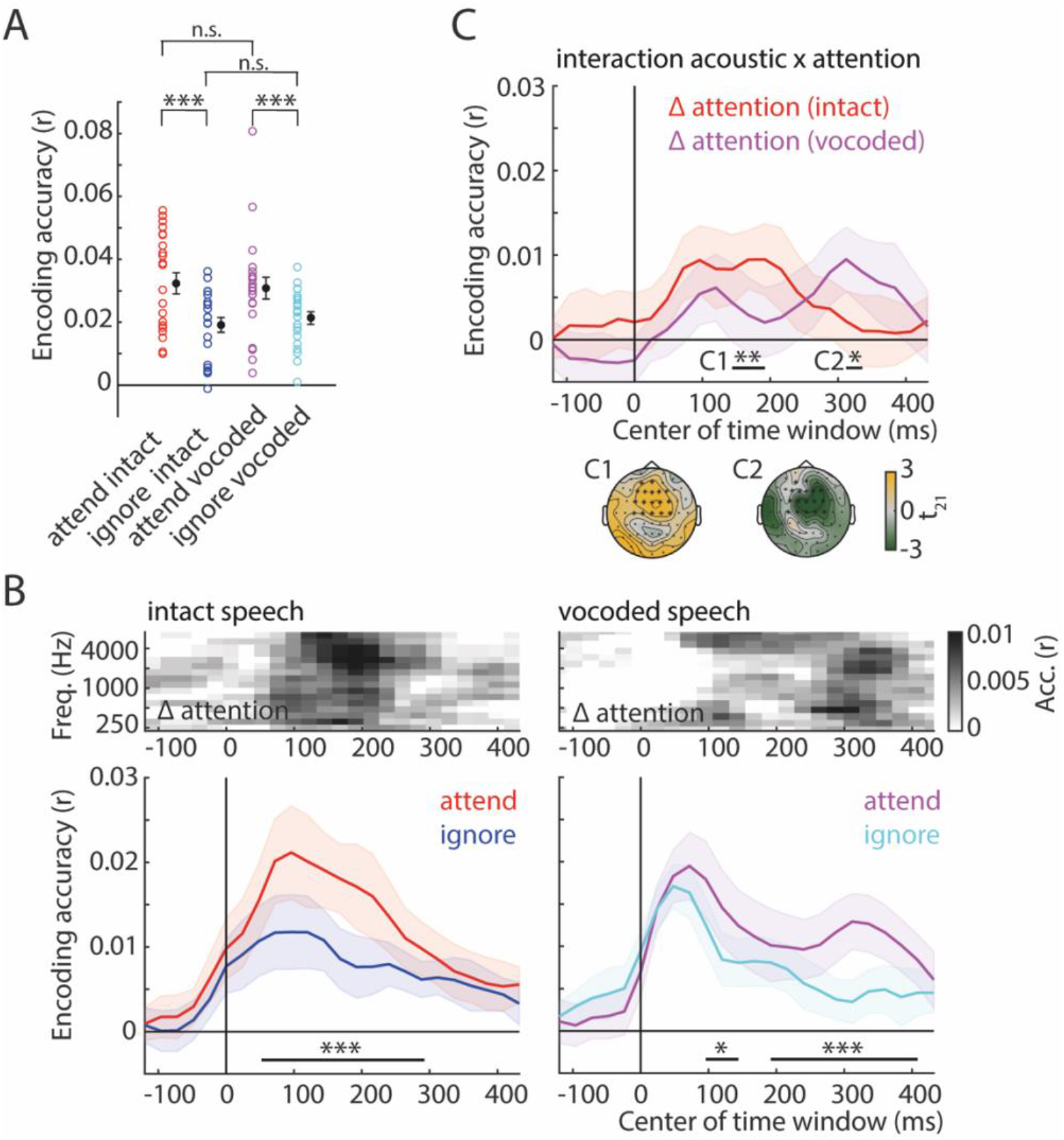
Acoustic degradation delays attentional enhancement of encoding accuracy. Encoding accuracy refers to the Pearson correlation of actual EEG data with the EEG data predicted by speech envelope models. (**A**) Encoding accuracy obtained when the entire sTRF (−150 to +450 ms) was used to predict EEG signals. Circles, dots, and error bars show single-subject, average encoding accuracy ±1 SEM, respectively. (**B**) Encoding accuracy obtained for sliding windows along the sTRF (window length: 48 ms; values on the x-axis correspond to centres of time windows). Solid lines and shaded areas indicate mean encoding accuracy and 95-% confidence intervals, respectively. Horizontal black lines indicate significant clusters for the difference of encoding accuracy for attended versus ignored speech. Time-frequency spectra on top show encoding accuracy for the effect of attention (Δ attention: attend–ignore). (**C**) In order to illustrate the attentional enhancement of encoding accuracy, data from (B) were contrasted for attended vs. ignored speech (Δ attention), separately for intact (red) and vocoded speech (purple). Horizontal black lines indicate significant clusters for the difference in the attentional enhancement for intact versus vocoded speech. Topographic maps show *t*-values for significant clusters; electrodes of significant clusters are highlighted in black. *** *p* < 0.001; ** *p* < 0.01; * *p* < 0.05; n.s. not significant.

First, when we used predicted EEG signals derived from the entire sTRF (Fig. 4A; −150 to +450 ms), encoding accuracy was significantly increased for attended compared to ignored intact speech (*p_permutation_* = 0.0002) and vocoded speech (*p_permutation_* = 0.0008). However, the acoustic detail x attention focus interaction was not significant (statistical comparison of the attended–ignored difference for intact versus vocoded speech; *p_permutation_* = 0.197), such that acoustic detail (intact vs. vocoded) had no significant effect on encoding accuracy for attended speech (*p_permutation_* = 0.66) or ignored speech (*p_permutation_* = 0.36). In other words, when predictions are based on the entire duration of the sTRF, attention enhances the encoding accuracy of intact and vocoded target speech to similar extents.

Next, we investigated the temporally-resolved dynamics of the attentional enhancement of intact versus vocoded speech (Fig. 4B). The attentional enhancement of encoding target versus distracting speech was signified by one cluster for intact speech (48-288 ms; *cluster p-value* = 0.0004; *t*-sum = 1,132; No. of cluster bins = 288) and by two clusters for vocoded speech (first cluster: 96-144 ms; *cluster p-value* = 0.039; *t*-sum = 64; No. of cluster bins = 24; second cluster: 192-408 ms; *cluster p-value* = 0.0002; *t*-sum = 521; No. of cluster bins = 168). The most important finding of this study resulted from the contrast of the attentional enhancement of encoding accuracy for intact versus vocoded speech (Fig. 4C): While the attentional enhancement was stronger for intact than vocoded speech in a relatively early cluster (144-192 ms, *cluster p-value* = 0.0054; *t*-sum = 94; No. of cluster bins = 34), the reverse pattern – stronger attentional enhancement for vocoded than intact speech – was observed in a later cluster (312-336 ms; *cluster p-value* = 0.0156; *t*-sum = −68; No. of cluster bins = 23).

It must be noted that cluster-based permutation tests do not establish significance of latencies of the observed clusters (Sassenhagen & Draschkow, 2019). In this sense, it is not warranted to draw inference about when in time the two observed acoustics x attention effects on encoding accuracy (in clusters C1 and C2 in Fig. 4C) started and ended being significant. However, since the two clusters observed here show significant effects in opposite directions, it can be excluded that the two underlying effects overlap in time (given that the topographical distribution of clusters is very similar). Thus, the attentional enhancement of encoding accuracy for vocoded speech is delayed compared to intact speech (see Supplementary Fig. S3 for an additional statistical analysis that confirms the latency difference).

## Discussion

The present study was designed to explore the neural processing constraints on selective attention in challenging listening situations imposed by unilateral spectral degradation, as is common in unilateral cochlear implant users. To this end, we studied neural tracking of the envelopes of two continuous, spatially separated speech signals. The most important results can be summarized as follows. First, there was a detrimental effect of unilateral spectral degradation on behaviour: Although high overall accuracy (~87%) indicates that participants mastered the listening task quite well, the proportion of correctly answered content questions about to-be-attended speech was about 10% reduced when it was spectrally degraded versus intact. Second, there was an effect of unilateral spectral degradation on neural responses: The preferential neural tracking of attended versus ignored speech was more pronounced for intact speech during earlier neural response components, but stronger for degraded speech for later components. These findings suggest that unilateral acoustic degradation impairs mainly the attentional selection of target speech against distraction by delaying it.

### Effects of unilateral acoustic degradation on tracking target versus distracting speech

Participants’ behavioural responses in the present study show a clear benefit of attending to intact compared to spectrally degraded speech on one side. The reason for this is likely a superposition of two effects. First, target speech comprehension is superior for speech that is spectrally less degraded (e.g. Obleser & Weisz, 2012; Shannon et al., 1995). Second, irrelevant speech distractors that are spectrally degraded interfere less with target speech reception (Westermann & Buchholz, 2017) and memory for target speech (e.g. Ellermeier et al., 2015; Senan et al., 2018; Wöstmann & Obleser, 2016). In this sense, the present study implemented one listening condition that was favourable on two accounts (attend to intact speech, ignore vocoded speech) and another listening condition that was unfavourable on two accounts (attend to vocoded speech, ignore intact speech; Supplementary Figure S2 shows the TRF grouped according to these two listening conditions). Note that these listening conditions correspond to rather extreme cases on the continuum of two-talker situations experienced in case of unilateral vocoding, such as they occur for unilateral cochlear implant (CI) users.

Although effects of bilateral spectral degradation of speech in multi-talker situations on neural envelope tracking have been investigated before (Ding et al., 2013; Kong et al., 2015; Rimmele et al., 2015), less is known about possible effects of unilateral spectral degradation. Here, we show that the typical components of the neural speech tracking response (P1, N1, P2) are present for unilateral intact as well as noise vocoded target speech, but that N1 and P2 were reduced for ignored speech distractors (Fig. 2 & 3). The general increase of the early P1 response component for noise-vocoded compared with intact speech (largely irrespective of attention) is most likely due to the fact that noise-vocoded speech is spectrally broader and thus elicits a more pronounced early neural response in auditory regions. This might also be related to the observation that noise-vocoded speech with fewer frequency channels is perceived as being louder (Wöstmann, Lim, et al., 2017).

Interestingly, a late negative N2 component signified the tracking of the attended speech envelope, particularly when it was noise vocoded. This finding somewhat differs from a previous study that presented listeners with spatially mixed competing speech signals (Fiedler et al., 2019), where we found a larger N2 component in the neural tracking of ignored speech (although with a shorter latency). Despite attempts to assign specific perceptual and cognitive processes to the different components of the neural speech tracking response (for review, see Brodbeck & Simon, 2020) the underlying mechanisms, particularly of the longer-latency components, are still unclear.

Importantly, the effect of unilateral vocoding on the neural tracking of attended versus ignored speech in the present study was best described as a temporal delay, instead of a reduced magnitude of attentional selection: Early neural tracking responses to intact unilateral speech, mainly during the N1 and P2 components, were larger for attended versus ignored speech. Attending versus ignoring vocoded speech increased the magnitude of later neural tracking responses, mainly during the P2 and N2 components. This finding is generally in line with a recent study that found delayed attentional modulation of neural speech tracking responses in bilateral CI users compared with normal-hearing controls (Paul et al., 2020).

What are the possible perceptual and cognitive processes associated with the observed temporally delayed attentional separation of attended versus ignored speech in case of unilateral acoustic degradation? One obvious interpretation is that in case of intact speech, the attentional selection is accomplished early, and largely in parallel with the formation of distinct auditory objects (Shinn-Cunningham, 2008). However, in the case of unilateral acoustic degradation, attentional selection happens later in time, potentially after the formation of competing auditory objects has been accomplished. Of note, neural responses to one speech signal might be significantly affected by the acoustic properties of the masking distractor in the present study. In this sense, an intact speech distractor that is highly intelligible increases the degree of informational masking, which might eventually result in delayed stream segregation (Ezzatian et al., 2015).

The late negative N2 TRF component observed in the present study is somewhat reminiscent of the Nd (negative difference) event-related potential (ERP) component that refers to a fronto-central scalp negativity in response to attended versus ignored tone stimuli (Hansen & Hillyard, 1980). The Nd component has also been observed in response to auditory probes in the attended versus ignored speech stream in multi-talker situations with competing speech signals presented at different spatial locations (Lambrecht et al., 2011; Münte et al., 2010; Nager et al., 2008). Although we can only speculate about the underlying cognitive mechanisms, we consider two processes that might give rise to the delayed attentional selection for vocoded speech in the time interval of the N2 component. First, it might be that perceptual restoration of vocoded target speech is more time consuming, such that the attentional separation of target and distractor is delayed in time. Second, it could be that participants use different strategies for the attentional selection of unilateral vocoded versus intact speech. For instance, it might be that the attentional selection of intact target speech exploits mainly the frequency separation of the competing speech signals, whereas the attentional selection of vocoded target speech is based on the additional use of spatial cues (for use of space and frequency cues in speech segregation, see Bonacci et al., 2020).

Of note, results of the present study emphasize the importance of a time-resolved analysis of the neural speech tracking response. In other words, a non time-resolved analysis on its own (such as the analysis of encoding accuracy in Fig. 4A) would have overlooked the temporally delayed attentional separation of unilateral vocoded target versus distractor speech. However, the spectrally-resolved analysis of speech tracking responses (sTRF), on the basis of speech envelopes extracted from individual frequency bands of the acoustic signal, did not reveal additional insights in the present study. That is, acoustics- and attention-induced modulations of the sTRF mostly covered a broad frequency range (1–8,000 Hz). It might be that the present study, that presented speech materials with envelopes correlated across neighbouring frequency bands and recorded the net EEG signal originating from large regions of underlying cortical tissue, lacked the required frequency-specificity. Future studies might thus utilize higher frequency-specificity on both, the level of the acoustic stimulus (e.g. by using synthesized stimuli with uncorrelated envelopes across frequency bands), and on the level of the neural response (e.g. through high-density cortical surface recordings in auditory regions; Hullett et al., 2016).

### What are the implications for listeners with a unilateral cochlear implant?

Although the present study was designed to simulate some aspects of hearing with a unilateral CI, the results should be transferred to actual unilateral CI user with caution. We presented normal-hearing listeners with unilateral vocoded versus intact speech without prior adaptation or speech comprehension training of spectrally degraded materials. It is thus conceivable that the observed delayed attentional selection of vocoded target speech against distraction is subject to change, such that its latency decreases in the course of rehabilitation of CI users (for a recent review on auditory cortical plasticity in CI users, see Glennon et al., 2020). Furthermore, the present study simulated hearing with a CI on one side with normal hearing on the contralateral side.

While such a scenario mimics hearing in people with single-sided deafness (SSD) after CI implantation, a significant amount of unilateral CI users suffers from hearing loss also on the contralateral side, which is treated with a hearing aid in case of bimodal listeners (for comparison of spatial speech-in-noise performance of SSD and bimodal listeners, see e.g. Williges et al., 2019).

Although dichotic listening is a useful experimental paradigm that achieves maximal spatial separation of competing sound sources, it must be noted that listening situations with a target signal present at 90° to one side under distraction from 90° to the other side are rare. Furthermore, the dichotic sound presentation over headphones in the present study did not allow for a partial mixture of the two competing signals before entering either ear, which would happen in real life listening situations with free field sound presentation. However, a pilot study with free field presentation of competing sound sources (see Supplementary Materials and Supplementary Fig. S1) revealed that unilateral vocoding in normal-hearing listeners and listening with a unilateral CI in a small group of CI users produced comparable speech comprehension detriments resulting from spectral degradation. Thus, it is conceivable that the present experimental paradigm simulates some challenges faced when listening with a unilateral CI. Nevertheless, to achieve higher ecological validity, future studies should use free field presentation and locate one sound source in the front and another one the left or right side, as we have done recently in normal-hearing listeners (Wöstmann et al., 2019). In theory, smaller spatial separation of sound sources should challenge the attentional separation of competing speech signals. Whether systematic variation in spatial separation of sound sources interacts with effects of spectral degradation on speech tracking is a question for future research.

Mechanistically, it is conceivable that the delayed attentional separation of unilateral vocoded target versus distractor speech constitutes a reason for compromised speech-in-noise comprehension in CI users. An ensuing question to be answered in future studies that implement single-trial speech comprehension measures, is whether the temporal delay of attentional selection in the speech tracking response relates to target speech comprehension. Alternatively, the delayed attentional separation might signify larger listening effort, which has been shown to increase with decreasing spectral resolution of speech (e.g. Pals et al., 2013).

## Conclusion

The present study presents evidence that unilateral acoustic degradation does not preclude the attention-induced neural selection of target speech from one side against concurrent distraction from the other. Instead, unilateral acoustic degradation delays this neural selection process. These findings improve our understanding of the neurophysiological basis of selective auditory attention to ongoing speech. Furthermore, we here identify a candidate neural signature of aggravated speech-in-noise comprehension under spectral degradation of the acoustic input, which has a potency in better explaining some of the listening challenges experienced by unilateral CI users.

## Acknowledgements

The authors acknowledge the financial support of Cochlear Inc. (Grant to JO and MW) and the European Research Council (Grant ERC-CoG-2014-646696 to JO). We thank Sebastian Puschmann and Martin Orf for their help with the mTRF toolbox.

## Data accessibility statement

The data are available from the corresponding author upon request.

## Supplementary materials

### Unilateral acoustic degradation simulates some aspects of hearing with a CI

In a pilot experiment, we directly compared the effects of unilateral speech vocoding in normal-hearing listeners (*N* = 11; age = 21–31 years; 9 females) to hearing with a Cochlear Implant (CI) in unilateral CI users (*N* = 6; age = 20–80 years; 5 females; CI-side = 3 left, 3 right). Five CI users had a hearing aid on the side contralateral to the CI (i.e., *bimodal CI users*).

In a cued auditory spatial attention task (Fig. S1A), participants were presented simultaneously with two spoken numbers, one presented from a loudspeaker positioned on the left (−90°) and the other presented from a loudspeaker positioned on the right (+90°). A visual cue in the beginning of each trial indicated the to-be-attended side (left/right). At the end of a trial, participants had to enter the number that appeared on the to-be-attended side on a keyboard. We used an adaptive tracking procedure to assess the speech reception threshold corresponding to the signal-to-noise ratio of the two spoken numbers necessary to arrive at 50% task accuracy (SRT_50_).

For normal-hearing listeners, lateralized speech signals presented on each trial were spectrally intact on one side and spectrally vocoded on the other side (using noise-vocoding with 16 bands for *N* = 6 and with 4 bands for *N* = 5). For CI users, all presented speech materials were intact. SRT_50_ values were determined for intact versus vocoded speech in normal-hearing listeners and for speech presented on the CI side versus non-CI side for CI users.

Figure S1 B&C show that unilateral vocoding with 16 bands in normal-hearing listeners did not differentially affect speech reception of vocoded speech (median SRT_50_ = −17.92 dB) compared with intact speech (median SRT_50_ = −17.96 dB; *p_permutation_* = 0.34). However, unilateral vocoding with 4 bands significantly increased the speech reception threshold for vocoded speech (median SRT_50_ = −11.89 dB) compared with intact speech (median SRT_50_ = −16.91 dB; *p_permutation_* < 0.001). Similarly, CI users exhibited increased speech reception thresholds on the CI side (median SRT_50_ = 6.70 dB) compared with the non-CI side (median SRT_50_ = 2.93 dB; statistical comparison after removal of one outlier: *p_permutaion_* < 0.001).

Of note, overall speech reception thresholds (averaged across intact and vocoded for normal-hearing listeners and across non-CI and CI side for CI users), were considerably higher in CI users compared with normal-hearing listeners in the 16-band vocoding group (*p_permutation_* = 0.003) and with normal-hearing listeners in the 4-band vocoding group (*p_permutation_* = 0.01).

In sum, the results of this pilot experiment show that the difference in speech reception thresholds is comparable for unilateral CI users (when contrasting the non-CI side with the CI side) and for normal-hearing listeners (when contrasting intact with vocoded speech with a low number of frequency bands). In this sense, the present pilot data justify the assumption that a spatial listening task with intact versus spectrally degraded speech presented on the left versus right side simulates some features of listening with a unilateral CI.

**Figure S1.**
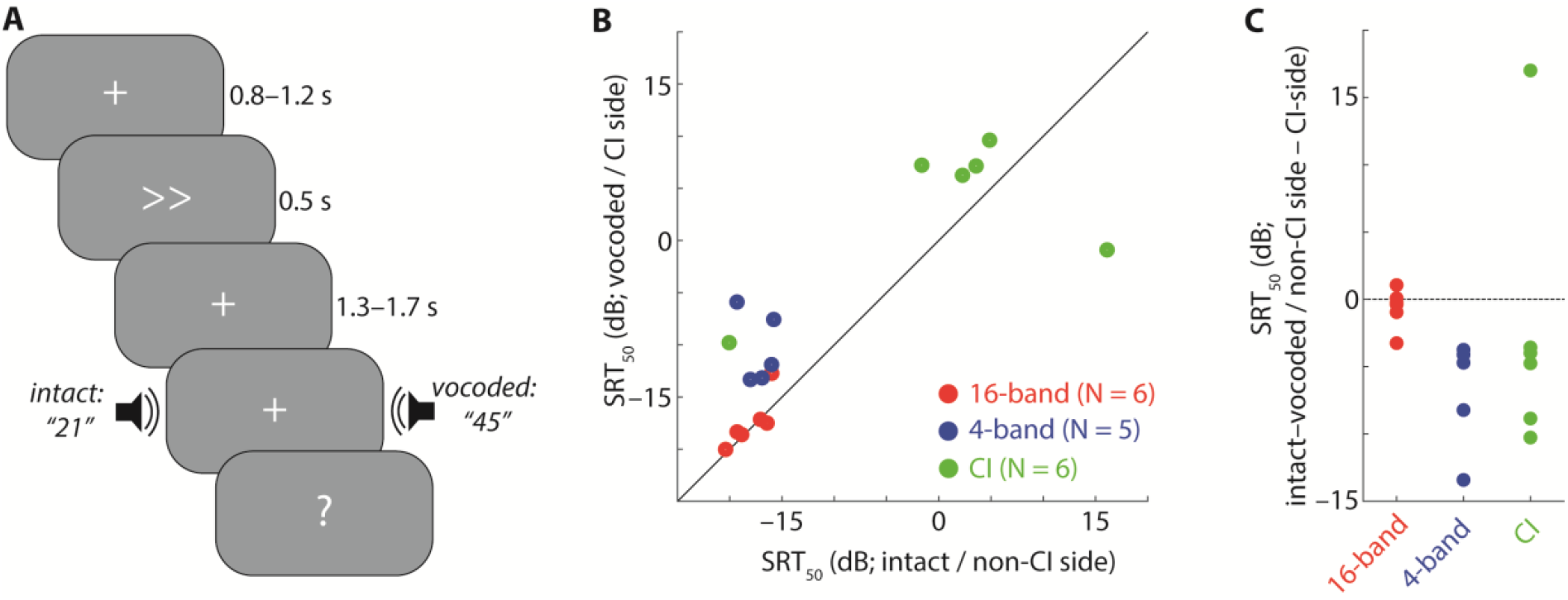
Experimental design and speech reception thresholds in a pilot experiment. (**A**) Design of spatial listening task. On each trial, a spatial cue guided participants’ attention to the left or right side. After a short delay, two spoken numbers were presented concurrently; one number was presented by the speaker on the left side and the other by the speaker on the right side. For normal-hearing participants, one of the numbers was spectrally degraded (using noise-vocoding). For unilateral CI-users, both numbers were intact, but one was presented on the CI-side and the other on the non-CI side. (**B**) 45-degree plot contrasts speech reception thresholds (SRT_50_) for attention to intact versus vocoded speech (using 16 or 4 vocoder bands) for normal-hearing participants, and for attention to the non-CI side versus CI side for unilateral CI-users. (**C**) Dots show single-subject SRTs for the difference intact–vocoded in case of normal hearing participants and for the difference non-CI side–CI side in case of unilateral CI users.

**Figure S2.**
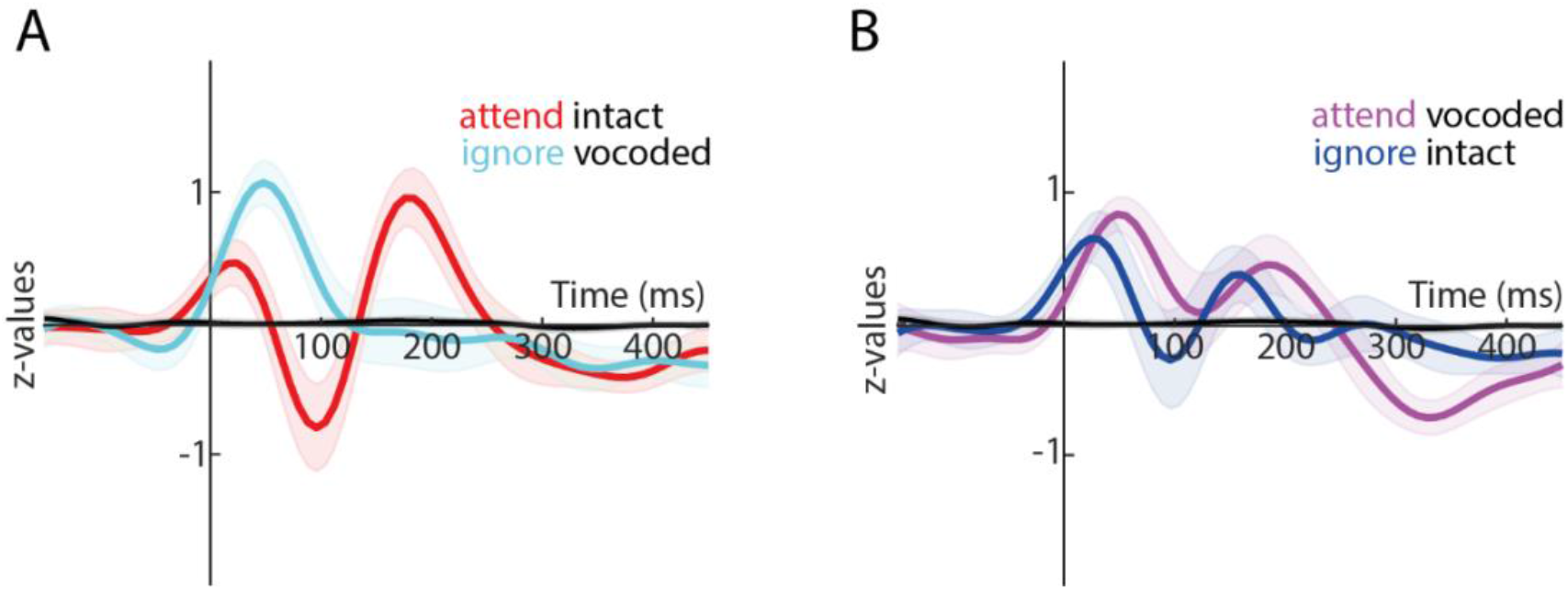
Temporal response functions grouped according to experimental settings. The temporal response function (TRF) was derived by averaging over nine central electrodes and frequencies of the sTRF. Shaded areas show 95-% confidence intervals. (**A**) Experimental setting 1: Participants attended to intact speech on one side while ignoring vocoded speech on the other side. (**B**) Experimental setting 2: Participants attended to vocoded speech on one side and ignored intact speech on the other side. Note that the data in this Figure are the same as in Figure 3D in the main article but grouped differently.

### Latency analysis of encoding accuracy modulation by acoustics and attention

In Figure 4C in the main article, we show that encoding accuracy for attended versus ignored speech was higher for intact speech in an earlier cluster but higher for vocoded speech in a later cluster. Here, we present results of an analysis that directly compares the latencies of these two effects. We implemented an analysis that avoids ambiguities associated with the comparison of peak latencies. For every participant, we first normalized the difference in encoding accuracy for attended–ignored speech between zero and one (Fig. S3 A). Next, we calculated the cumulative sum of normalized encoding accuracy across time lags −150 to +450 ms. For every participant and acoustic condition, we then determined for each percentage between 1 and 100 at which latency this percentage of the cumulative sum was reached. For statistical analysis we compared latencies for intact versus vocoded speech for percentages 1–100 with multiple dependent-samples *t*-tests (uncorrected). The result of this analysis demonstrate that the attentional enhancement of encoding accuracy occurred significantly earlier for intact than vocoded speech (Fig. S3 B). In more detail, latencies for which ~45–90% of the total encoding accuracy difference for attended versus ignored speech was reached were significantly longer for vocoded than intact speech.

**Figure S3.**
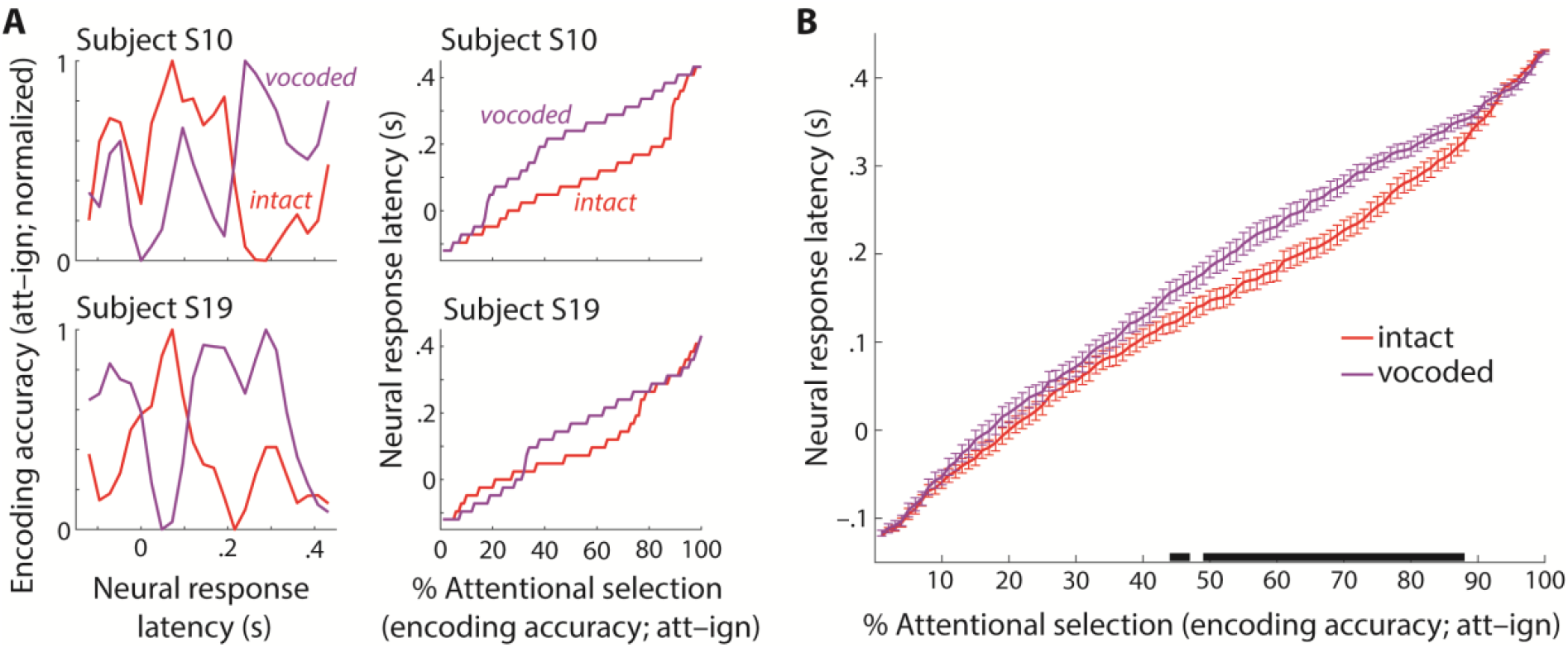
Latency differences of encoding accuracy modulation by acoustics and attention. (**A**) Calculation of the latency of attentional selection for two exemplary subjects. Left column: Normalized difference in encoding accuracy (attended–ignored) for intact (red) and vocoded speech (purple). Right column: Neural response latency as a function of the percentage of attentional selection (see text for details). (**B**) Solid lines and error bars show mean latency ±1 SEM (for the whole sample of *N* = 22 participants), respectively. The horizontal black bar indicates significant latency differences for intact versus vocoded speech (multiple dependent-sample *t*-tests; *p* < .05; uncorrected).

